## Supplemental material for "DMEM and EMEM are suitable surrogate media to mimic host environment and expand leptospiral pathogenesis studies using *in vitro* tools"

**S1 Table**. Top 10 up and downregulated leptospiral genes identified in whole blood (WB) compared to DMEM, EMEM, and HAN versus EMJH. p-value<0.01 and log2FC ± 2.

| | LIC # | Gene ID | WB | DMEM | EMEM | HAN |
|---|---|---|---|---|---|---|
| Upregulated genes | LIC_RS21185 | LIC_RS21185 | 11.88 | 9.96 | 10.20 | 8.66 |
| | LIC12890 | LIC_RS14805 | 8.72 | 6.35 | 6.15 | 6.25 |
| | LIC10109 | LIC_RS00560 | 7.28 | 6.54 | 6.90 | 6.88 |
| | LIC_RS20530 | LIC_RS20530 | 7.28 | 5.39 | 5.99 | - |
| | LIC12890 | LIC_RS14850 | 7.28 | 6.65 | 6.70 | 6.60 |
| | LIC13040 | LIC_RS15640 | 7.00 | 4.04 | 3.71 | 2.00 |
| | LIC_RS14820 | LIC_RS14820 | 6.99 | 3.86 | 3.70 | - |
| | LIC12631 | LIC_RS13475 | 6.63 | 5.86 | 7.40 | 5.84 |
| | LIC10108 | LIC_RS00555 | 6.29 | - | - | - |
| | LIC10462 | LIC_RS02405 | 6.28 | 5.86 | 7.52 | - |
| Downregulated genes | LIC13151 | LIC_RS16215 | -7.02 | -3.85 | -3.37 | -4.12 |
| | LIC12524 | LIC_RS12925 | -7.21 | -3.36 | -3.72 | -2.41 |
| | LIC10705 | LIC_RS03630 | -7.27 | -8.18 | -7.91 | - |
| | LIC10821 | LIC_RS04235 | -7.58 | -2.67 | -2.61 | - |
| | LIC12988 | LIC_RS15375 | -7.65 | -3.62 | -3.09 | - |
| | LIC10708 | LIC_RS03645 | -7.80 | -5.77 | -5.21 | - |
| | LIC10704 | LIC_RS03625 | -8.73 | -9.13 | -8.06 | -2.38 |
| | LIC_RS02170 | LIC_RS02170 | -9.06 | - | - | - |
| | LIC10707 | LIC_RS03640 | -9.97 | - | - | - |
| | LIC_RS23399 | LIC_RS23400 | -18.24 | - | - | - |

**S2 Table.** Biological pathways identified with genes differentially expressed between different growth conditions and environments compared to EMJH (*P* <0.05).

| **Biological Pathways** | **DMEM** | **EMEM** | **HAN** | **WB** |
|---|---|---|---|---|
| Cob(II)yrinate a,c-diamide biosynthesis I (early cobalt insertion)* | 0.001 | 0.000 | 0.001 | 0.033 |
| Cobamide biosynthesis* | - | 0.001 | 0.001 | 0.005 |
| Fatty acid and lipid degradation# | 0.001 | 0.032 | - | 0.029 |
| Fatty acid and lipid biosynthesis# | 0.030 | 0.048 | 0.008 | - |
| Sphingomyelin metabolism | 0.012 | 0.045 | 0.012 | - |
| Thiamine biosynthesis | 0.035 | - | - | - |
| S-adenosyl-L-methionine biosynthesis | - | 0.002 | - | - |
| L-methionine degradation | - | 0.016 | - | - |
| Metabolic clusters | - | - | - | 0.032 |
| 2-Deoxy-D-ribose degradation | - | - | - | 0.038 |
| TCA cycle I (prokaryotic) | - | - | - | 0.039 |
| tRNA charging | - | - | - | 0.042 |
| Enzyme cofactor biosynthesis | 0.000 | 0.000 | 0.004 | - |

*Upregulated pathways, #Downregulated pathways.

**S3 Table**. Biological pathways identified with genes differentially expressed between DMEM and EMJH media.

| **Pathways** | ***P*** | **Matches** |
|---|---|---|
| Enzyme cofactor biosynthesis | $1.57 \times 10^{-04}$ | LIC_RS18015 // LIC_RS18625 // LIC_RS18615 // LIC_RS18600 // LIC_RS18610 // LIC_RS13725 // LIC_RS18650 // LIC_RS18630 // LIC_RS18635 // LIC_RS18620 // LIC_RS04135 |
| cob(II)yrinate a,c-diamide biosynthesis I (early cobalt insertion) | $8.09 \times 10^{-04}$ | LIC_RS18650 // LIC_RS18630 // LIC_RS18635 // LIC_RS18620 |
| Cobamide biosynthesis | $8.09 \times 10^{-04}$ | LIC_RS18625 // LIC_RS18615 // LIC_RS18600 // LIC_RS18610 |
| Fatty acid and lipid degradation | 0.0015 | LIC_RS00485 // LIC_RS05035 // LIC_RS16980 // LIC_RS16975 // LIC_RS07230 |
| Sphingomyelin metabolism | 0.015 | LIC_RS13475 // LIC_RS13480 |
| Fatty acid and lipid biosynthesis | 0.030 | LIC_RS07230 // LIC_RS11880 // LIC_RS17080 // LIC_RS13480 // LIC_RS13475 // LIC_RS00485 |
| Thiamine biosynthesis | 0.035 | LIC_RS13725 // LIC_RS04135 |

**S4 Table**. Biological pathways identified with differentially expressed genes between EMEM and EMJH media.

| **Pathways** | ***P*** | **Matches** |
|---|---|---|
| Cob(II)yrinate a,c-diamide biosynthesis I (early cobalt insertion) | $8.09 \times 10^{-05}$ | LIC_RS18650 // LIC_RS18660 // LIC_RS18630 // LIC_RS18635 // LIC_RS18620 // LIC_RS18655 |
| Enzyme cofactor biosynthesis | $8.80 \times 10^{-05}$ | LIC_RS18015 // LIC_RS17280 // LIC_RS18625 // LIC_RS18615 // LIC_RS18600 // LIC_RS18610 // LIC_RS18420 // LIC_RS18410 // LIC_RS06950 // LIC_RS13725 // LIC_RS18650 // LIC_RS18660 // LIC_RS18630 // LIC_RS18635 // LIC_RS18620 // LIC_RS18655 // LIC_RS04135 |
| Cobamide biosynthesis | 0.0012 | LIC_RS17280 // LIC_RS18625 // LIC_RS18615 // LIC_RS18600 // LIC_RS18610 |
| S-adenosyl-L-methionine biosynthesis | 0.0020 | LIC_RS18420 // LIC_RS18410 // LIC_RS06950 |
| L-methionine degradation | 0.016 | LIC_RS18410 // LIC_RS06950 |
| Fatty acid and lipid degradation | 0.032 | LIC_RS05035 // LIC_RS16980 // LIC_RS16975 // LIC_RS07230 // LIC_RS18395 |
| Sphingomyelin metabolism | 0.045 | LIC_RS13475 // LIC_RS13480 |
| Fatty acid and lipid biosynthesis | 0.048 | LIC_RS07230 // LIC_RS18395 // LIC_RS07775 // LIC_RS11880 // LIC_RS17080 // LIC_RS18310 // LIC_RS13475 // LIC_RS13480// LIC_RS07310 |

**S5 Table**. Biological pathways identified with differentially expressed genes between HAN and EMJH media.

| **Pathways** | ***P*** | **Matches** |
|---|---|---|
| Cob(II)yrinate a,c-diamide biosynthesis I (early cobalt insertion) | $8.09 \times 10^{-04}$ | LIC_RS18650 // LIC_RS18630 // LIC_RS18635 // LIC_RS18620 |
| Cobamide biosynthesis | $8.09 \times 10^{-04}$ | LIC_RS18625 // LIC_RS18615 // LIC_RS18600 // LIC_RS18610 |
| Enzyme cofactor biosynthesis | 0.0037 | LIC_RS18625 // LIC_RS18615 // LIC_RS18600 // LIC_RS18610 // LIC_RS13725 // LIC_RS18650 // LIC_RS18630 // LIC_RS18635 // LIC_RS18620 |
| Fatty acid and lipid biosynthesis | 0.0077 | LIC_RS07230 // LIC_RS11880 // LIC_RS17080 // LIC_RS10540 // LIC_RS13475 // LIC_RS13480 // LIC_RS07310 |
| Sphingomyelin metabolism | 0.012 | LIC_RS13475 // LIC_RS13480 |

**S6 Table**. Biological pathways identified with differentially expressed genes between WB and EMJH.

| **Pathways** | ***P*** | **Matches** |
|---|---|---|
| Cobamide biosynthesis | $4.83 \times 10^{-03}$ | LIC_RS18625 // LIC_RS07865 // LIC_RS18615 // LIC_RS18600 // LIC_RS18610 |
| Fatty acid and lipid degradation | $2.89 \times 10^{-02}$ | LIC_RS00485 // LIC_RS05035 // LIC_RS16980 // LIC_RS02070 // LIC_RS16975 // LIC_RS07230 |
| Metabolic clusters | 0.032 | LIC_RS16665 // LIC_RS17820 // LIC_RS13150 // LIC_RS11615 // LIC_RS01810 // LIC_RS04700 // LIC_RS16635 // LIC_RS04695 // LIC_RS08130 // LIC_RS07755 // LIC_RS16075 // LIC_RS13280 // LIC_RS10010 // LIC_RS13155 // LIC_RS02230 // LIC_RS12605 // LIC_RS00495 |
| Cob(II)yrinate a,c-diamide biosynthesis I (early cobalt insertion) | 0.033 | LIC_RS18650 // LIC_RS18630 // LIC_RS18635 // LIC_RS18620 |
| 2-Deoxy-D-ribose degradation | 0.038 | LIC_RS05035 // LIC_RS16980 // LIC_RS02070 |
| TCA cycle I (prokaryotic) | 0.039 | LIC_RS10225 // LIC_RS09090 // LIC_RS15025 // LIC_RS13190 // LIC_RS13185 |
| tRNA charging | 0.042 | LIC_RS17820 // LIC_RS13150 // LIC_RS11615 // LIC_RS01810 // LIC_RS04700 // LIC_RS16635 // LIC_RS04695 // LIC_RS08130 // LIC_RS07755 // LIC_RS16075 // LIC_RS13280 // LIC_RS10010 // LIC_RS13155 // LIC_RS02230 // LIC_RS12605 |

**S7 Table**. Percentual of differently expressed genes between different media conditions and dialysis membrane chamber (DMC) compared to whole blood (WB) samples.

| Media | Up-regulated genes | | | Down-regulated genes | | | % of all WB differentially regulated genes |
|---|---|---|---|---|---|---|---|
| | n | Shared with WB (%) | % of all WB up regulated genes | n | Shared with WB (%) | % of all WB down regulated genes | |
| **DMEM** | 173 | 136 (79) | 35 | 241 | 156 (65) | 45 | 40 |
| **EMEM** | 241 | 174 (72) | 44 | 264 | 171 (65) | 50 | 47 |
| **HAN** | 167 | 92 (55) | 23 | 104 | 58 (50) | 17 | 20 |
| **DMC*** | 110 | 47 (43) | 12 | 56 | 19 (34) | 6 | 9 |

* Caimano et al, PLoS Pathogens 2014 Mar 13;10(3):e1004004

**S8 Table**. Differential expression of genes related to lipid A biosynthesis in each medium compared to EMJH

| Gene | ID | DMEM | | EMEM | | HAN | | WB | |
|---|---|---|---|---|---|---|---|---|---|
| | | **log2FC** | **FDR** | **log2FC** | **FDR** | **log2FC** | **FDR** | **log2FC** | **FDR** |
| *lpxC* | | 0.57 | $9.5 \times 10^{-06}$ | 0.9 | $1.1 \times 10^{-12}$ | 1 | $1.0 \times 10^{-16}$ | 2.42 | $1 \times 10^{-77}$ |
| *lpxK* | LIC12579 | 1.42 | $3.3 \times 10^{-20}$ | 1.37 | $7.2\times 10^{-19}$ | 0.41 | $1.1 \times 10^{-02}$ | 1.6 | $1 \times 10^{-08}$ |
| *lpxH* | LIC20219 | - | - | - | - | - | - | 0.75 | $2 \times 10^{-02}$ |
| *lpxD1* | LIC13046 | 0.36 | $5.0 \times 10^{-02}$ | 1.19 | $8.9 \times 10^{-11}$ | 0.67 | $3.7 \times 10^{-04}$ | 0.69 | $4 \times 10^{-04}$ |
| *lpxB* | LIC12579 | 0.6 | $4.8 \times 10^{-07}$ | 0.5 | $2.6 \times 10^{-05}$ | 0.25 | $4.6 \times 10^{-02}$ | 0.49 | $2 \times 10^{-03}$ |
| *lpxA* | | -0.82 | $7.5 \times 10^{-13}$ | -0.5 | $1.5 \times 10^{-05}$ | -0.91 | $1.2 \times 10^{-15}$ | -1.15 | $4\times 10^{-13}$ |
| *lpxI* | LIC12578 | - | - | - | - | -0.28 | $1.6 \times 10^{-02}$ | -1.18 | $2 \times 10^{-07}$ |
| *lpxD2* | LIC13469 | -0.68 | $1.9 \times 10^{-04}$ | - | - | - | - | - | - |
